# Environmentally Relevant Lead Exposure Alters Cell Morphology and Expression of Neural Hallmarks During SH-SY5Y Neuronal Differentiation

**DOI:** 10.1101/2025.02.17.638689

**Authors:** Rachel K. Morgan, Anagha Tapaswi, Katelyn M. Polemi, Jenna L. Miller, Jonathan Sexton, Kelly M. Bakulski, Laurie K. Svoboda, Dana C. Dolinoy, Justin A. Colacino

**Affiliations:** Department of Environmental Health Sciences, School of Public Health, University of Michigan, Ann Arbor, MI 48109, USA; Department of Medicinal Chemistry, College of Pharmacy, University of Michigan, Ann Arbor, MI 48109, USA; Department of Epidemiology, School of Public Health, University of Michigan, Ann Arbor, MI 48109, USA; Department of Pharmacology, University of Michigan Medical School, Ann Arbor, MI 48109, USA; Department of Nutritional Sciences, School of Public Health, University of Michigan, Ann Arbor, MI 48109, USA; Program in the Environment, University of Michigan College of Literature, Sciences, and the Arts and the School of Environment and Sustainability, Ann Arbor, MI 48109

**Keywords:** neurotoxicology, neurodevelopment, morphology, neural differentiation, lead

## Abstract

Lead (Pb) continues to be a public health burden, in the US and around the world, and yet the effects of historical and current exposure levels on neurogenesis are not fully understood. Here we examine the effects of a range of environmentally relevant Pb concentrations (0.16μM, 1.26μM, and 10μM Pb) relative to control on neural differentiation in the SH-SY5Y cell model. Pb exposure began on Day 5 and continued throughout differentiation at Day 18. We assessed morphological measures related to neurogenesis at several time points during this process, including the expression of proteins key in neural differentiation (β-tubulin III and GAP43), cell number and size, as well as the development of neurites. The bulk of detectable changes occurred with 10μM Pb exposure, most notably that of β-tubulin III and GAP43 expression. Effects with the 0.16μM and 1.26μM Pb exposure conditions increased as differentiation progressed, with significant reductions in cell and nuclear size as well as the number and length of neural projections by Day 18. Best benchmark concentration (BMC) analysis revealed many of these metrics to be susceptible to levels of Pb at or below historically relevant levels. This work highlights the disruption of neurite formation and protein expression as potential new mechanisms by which environmentally relevant Pb exposure impacts neurogenesis and morphology and perturb cognitive health throughout the life course.

## Introduction

Despite major public health policy during the 20^th^ century, lead (Pb) exposure continues to be a persistent public health burden in the US and around the globe (Levin et al., 2021). In the US, it’s estimated that 1.28 million children under the age of 5 have blood lead levels (BLLs) greater than 3.5μg/dL (Ruckart et al., 2021) and worldwide, 800 million children have BLLs exceeding 5μg/dL (UNICEF, 2020). Developmental Pb exposure is a greater burden in low- and middle-income countries (Larsen & Sánchez-Triana, 2023) and remains on the World Health Organization’s list of ten chemicals of major public health concern (WHO, 2020). In the US, roughly half of the total population is thought to have been exposed to levels of Pb above the CDC’s current actionable level at some point during development (McFarland et al., 2022), meaning a significant proportion of adults today may be experiencing health outcomes related to exposures they experienced decades ago. Developmental exposure to Pb is associated with impairments to learning and behavior (Hou et al., 2013) and these effects are thought to be life-long (Glass et al., 2009; White et al., 1993). As efforts to reduce childhood and developmental Pb exposure and its effects continue in the US and around the world, there is a need for a better understanding of how early life Pb exposure drives cognitive impairment that lasts a lifetime, in order to better address the health needs of individuals exposed prior to stricter legislative measures.

One possible avenue by which early life Pb exposure may contribute to perturbed cognitive function during adulthood is by impacting neural stem cell differentiation. Neural stem cells are most prevalent during early life, when the majority of neurogenesis happens (Noctor et al., 2004), though neural stem cell function has been documented in several areas of the brain throughout the life course (Seri et al., 2001). During neural cell maturation, differentiating neurons rapidly develop axon and dendritic processes (Ribak et al., 2004; Silver et al., 1982), which is largely orchestrated through cytoskeletal arrangements including actin and microtubules (Forscher & Smith, 1988). Perturbations to synaptic function, via aberrant axon and dendritic development (H. Li et al., 2020), during neurogenesis is linked to adverse cognitive outcomes including learning and memory (Batool et al., 2019; Gruart et al., 2006), and dysregulation of microtubule function is associated with neurodegenerative disease (Calogero et al., 2019). In addition to cytoskeletal proteins, stemness factors such as GAP43 drive neurogenesis (Shen et al., 2004) via the regulation of intracellular signaling pathways (Mishra et al., 2008). Disruption of GAP43 expression and function is also linked to changes in learning and memory associated with age (Mingaud et al., 2008), and it may be that toxicant-induced perturbation to this protein may accelerate these effects. Indeed, Pb disrupts each of these processes essential for proper neural maturation. For example, 3-6μM Pb (equivalent to 62-124μg/dL Pb) disrupts cytoskeletal development and that of GAP43 expression in cell models of neurogenesis (Ayyalasomayajula et al., 2020; Scortegagna et al., 1998). Even lower concentrations of Pb, 0.07-.24μM Pb (1.45-5μg/dL Pb), perturb neurite (e.g., axon and dendrite) growth in developing neurons (Xie et al., 2023), demonstrating that more environmentally relevant levels of Pb exposure also perturb neurogenesis. However, it remains to be seen whether such levels perturb neural differentiation due to the disruption of stemness factors and cytoskeletal development and if these effects are differentiation-stage specific.

Here, we developed a high content imaging-based approach to evaluate the effects of environmentally relevant and continuous Pb exposure during neural differentiation on the expression of GAP43 and β-tubulin III, a major neural cytoskeletal protein in the SH-SY5Y neural stem cell model (Shipley et al., 2016). In conjunction with this, we also assessed the effects of exposure on neurite outgrowth, as quantified by the number of neural projections as well as their total length. We measured these outcomes at several points during differentiation to assess whether effects were stage-specific in an attempt to better understand how Pb exposure disrupts neurogenesis. These experiments build upon previous work which has found environmentally relevant levels of developmental Pb to perturb gene expression regulation in the adult mouse model (Morgan, Wang, et al., 2024; Petroff et al., 2023) as well as gene expression and pathways related to neurological function and neurodegenerative disease in cell models (Morgan, Tapaswi, et al., 2024). Given our previous results and those of the existing literature, we tested the hypothesis that environmentally relevant doses of Pb would suppress GAP43 and β-tubulin III expression, as well as that of neurite growth, in a dose-dependent manner and that these effects would intensify as neural differentiation progressed.

## Materials and Methods

### SH-SY5Y Differentiation and Exposure Conditions

SH-SY5Y cells (ATCC, Cat. #CRL-2266) were expanded according to manufacturer protocols in a 1:1 mixture of Eagle’s Minimum Essential Media (ATCC, Cat. #30-2003) and F-12 (Thermo, Cat. #11765-054), supplemented with 10% heat-inactivated fetal bovine serum (hiFBS) (Thermo, Cat. #A3840001) and 1% antibiotics (Thermo, Cat. #15140-122). SH-SY5Y cells were differentiated into neuron-like cells using previously published methods (Shipley et al., 2016). Briefly, on day (D) 0, cells were seeded at a density of 10,000 cells per well in a 24-well plate (Corning, Cat. #353047) and allowed to attach for 24 hours. Beginning on D1, cells were maintained for one week in serum-deprivation conditions (2.5% hiFBS) and 10μM retinoic acid (RA) (Sigma, Cat. #R2625). On D7, cells were split 1:1 onto new 24-well plates and maintained for an additional three days in 1% hiFBS and 10μM RA. Cells were split once more on D10 1:1 onto extracellular matrix (ECM) (Sigma, Cat. #E0282) coated plates and maintained in hiFBS-free media containing neurobasal medium (Thermo, Cat. #21103049), brain-derived neurotrophic factor (BDNF) (Thermo, Cat. #10908-010), and dibutyryl cyclic AMP (db-cAMP) (Fisher, Cat. #16980-89). Cells were considered fully differentiated into neuron-like cells on D18. A full description of media conditions can be found in **Table S1**.

Beginning on D5, differentiating SH-SY5Y cells were exposed to environmentally relevant concentrations of aqueous Pb acetate trihydrate (henceforth referred to as Pb) (Sigma, Cat. #316512) or control conditions. SH-SY5Y cells were allowed to begin differentiating in the absence of Pb to reduce stress and optimize cell viability. However, once Pb exposure began it was continuous for the remainder of the 18-day protocol. Exposure conditions included 0μM Pb (control), 0.16μM Pb, 1.26μM Pb, and 10μM Pb. These exposure conditions were chosen as they reflect current and historical exposures, as quantified via blood lead levels (BLLs). 0.16μM Pb corresponds to the blood lead reference value, as of 2024, set by the CDC (3.5μg/dL), which is based on the 97.5^th^ percentile of BLLs in children 1-5 years old in the US. 1.26μM Pb corresponds to BLLs prevalent in the US prior to the phase out of leaded gasoline and Pb-based paint in the 1970s (∼26μg/dL) (McFarland et al., 2022). 10μM Pb corresponds to BLLs greater than 200μg/dL, well above BLLs considered to be tolerable in humans (Markowitz, 2000). Unit conversion for μM to μg/dL was calculated using the molecular weight of Pb (207.2g/mol) and the equation μM Pb = μg Pb/((0.1dL/L)/(207.2g/mol)). Pb solution was added to cells with each media change (i.e., every 2-3 days). A full illustration of the differentiation and exposure protocol is provided in **Figure 1**.

**Figure 1:**
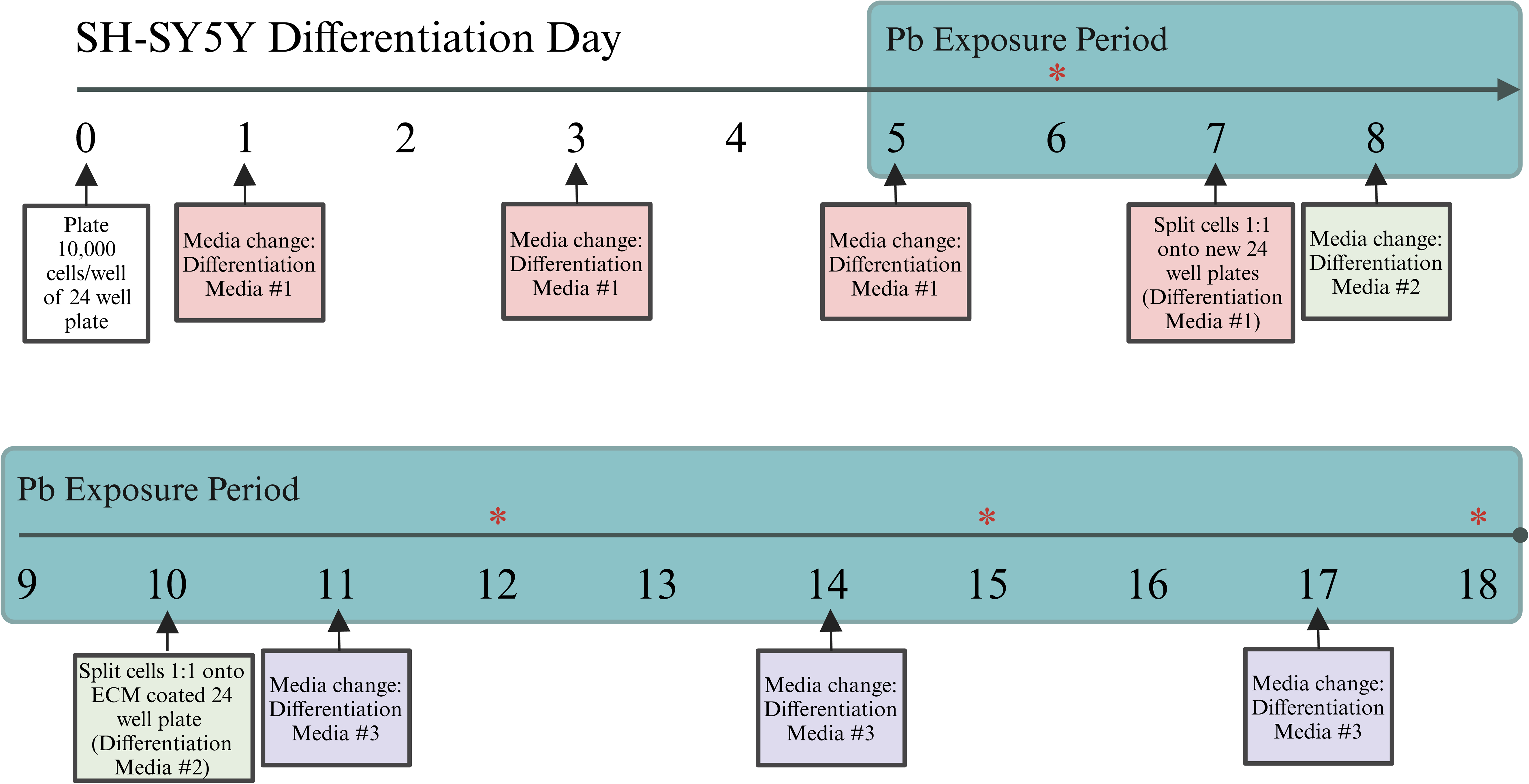
Experimental Workflow. SH-SY5Y cells were differentiated in the presence of 10uM retinoic acid (RA) for 18 days, with lead (Pb) exposure beginning on Day 5. Cells were collected on Days 6, 12, 15, and 18 (indicated by *) to assess changes in cellular morphology and the expression of neural proteins. A complete description of media recipes used can be found in Table S1. ECM: extracellular matrix.

**Figure 2:**
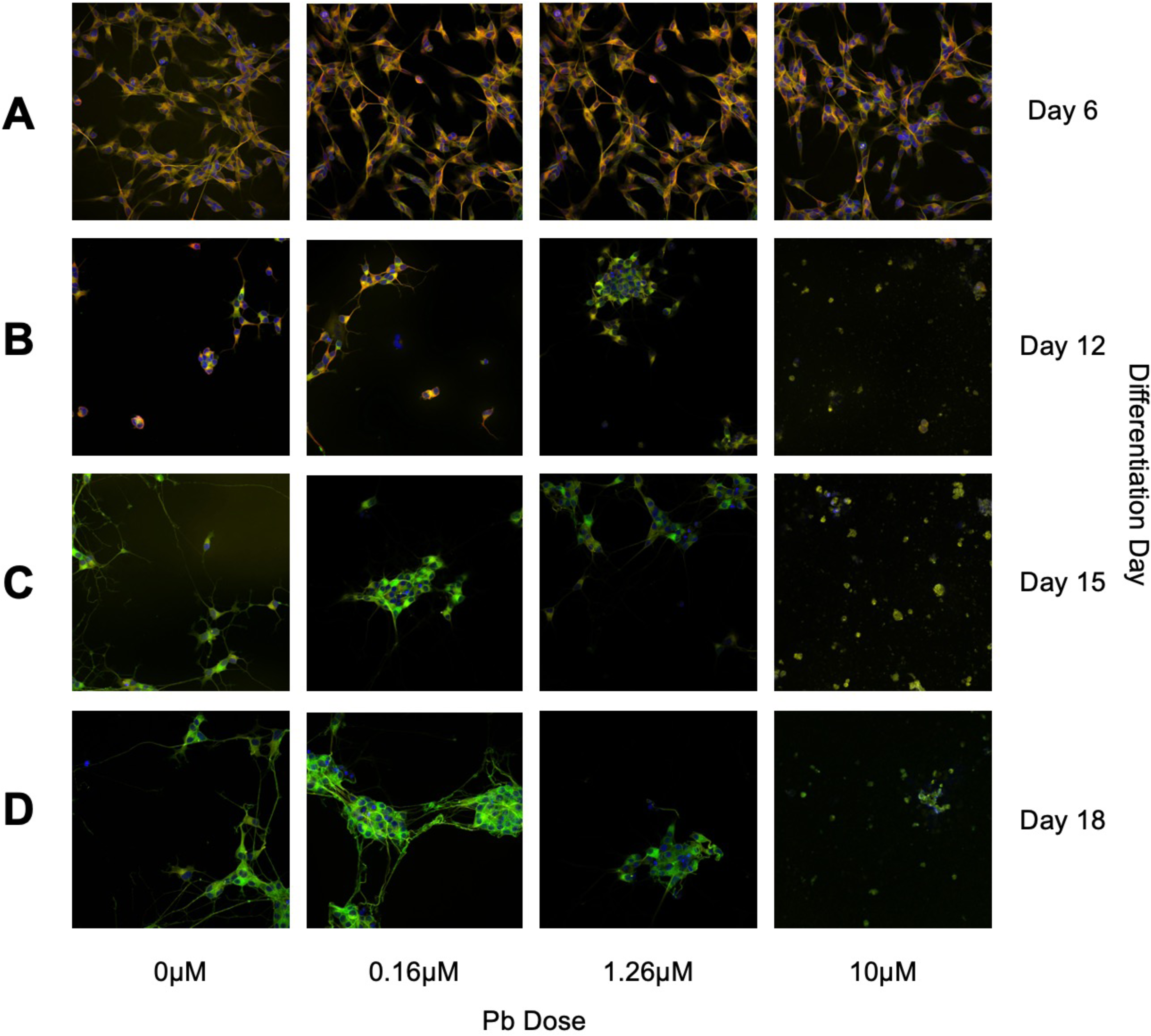
Representative Images of Cell Imaging. Cells were differentiated over the course of 18 days using methods adapted from Shipley et al., 2016. Beginning on Day 5 cells were exposed to a range of lead (Pb) concentrations (0uM, 0.16uM, 1.26uM, or 10uM Pb). Fixed and labeled cells were imaged using the Yokogawa CV8000 at 40X resolution. Cells were labeled using the nuclear stain Hoechst 33342, and immunofluorescently tagged antibodies for B-tubulin III and GAP43. Cells were fixed and stained on Days 6 (A), 12 (B), 15 (C), and 18 (D).

Six replicates for each exposure condition (n = 4) at each time point (n = 4) were included for analysis. Replicates refer to wells of a 24-well plate, in which each row of 6 wells corresponds to one concentration condition and one 24-well plate corresponds to one time point.

### Immunofluorescence Staining and Imaging

At each timepoint (D6, D12, D15, D18), SH-SY5Y cells were fixed and incubated with fluorescently-tagged antibodies for β-tubulin III and GAP43, and the nuclear stain Hoechst, in accordance with previously published protocols and manufacturer recommendations. β-tubulin III and GAP43 are markers of neuronal differentiation previously validated in SH-SY5Y-derived neuron-like cells (Shipley et al., 2016). β-tubulin III localizes throughout axonal and dendritic growth during neural differentiation (Guo et al., 2011), whereas GAP43 localizes to cell body-adjacent axon regions and the soma (Morita & Miyata, 2013). Hoechst binds to AT-rich regions of double stranded DNA, thus providing a reliable stain of the nucleus (Chazotte, 2011). Cells were fixed with 4% paraformaldehyde (PFA) solution (Thermo, Cat. #AA433689M) diluted in basic growth media (**Table S1**) and incubated at room temperature for 20 minutes, protected from light. PFA was removed and cells were rinsed once with a 1% solution of Tween 20 (Thermo, Cat. #85113) diluted in PBS, pH = 7.4 (Thermo, Cat. #10010023). Cells were then permeabilized with a 1% solution of Triton X (Thermo, Cat. #85111) diluted in PBS, incubated at room temperature for 15 minutes. Triton X solution was removed, and cells were rinsed twice with a 1% Tween 20 solution. Primary staining consisted of Hoechst 33342 (Hoechst, dilution 1:2000) (Thermo, Cat. #H3570), anti-β-tubulin III (mouse, dilution 1:150) conjugate antibody (eFluor660) (Thermo, Cat. #50-4510-82), and anti-GAP43 (rabbit, dilution 1:200) antibody (Abcam, Cat. #ab75810) in normal goat serum (NGS) (Thermo, Cat. #50062Z). Primary staining solution was applied to cells and incubated at room temperature protected from light for 1 hour. Primary staining solution was removed, and cells were rinsed three times with 1% Tween 20 solution. Secondary staining solution contained anti-rabbit (dil. 1:333), Alexa Fluor488) antibody (Abcam, Cat. #150077) diluted in NGS. Secondary staining solution was added to cells and incubated at room temperature protected from light for 1 hour. Secondary stain was removed, and cells were rinsed three times with 1% Tween 20 solution. Cells were stored in 1% Tween 20 solution at 4°C until imaging. A full description of SH-SY5Y fixation and labeling is included in **Supplementary File 1**.

Cells were imaged using the Yokogawa Cell Voyager (CV8000) microscope. Automated plate imaging was performed on the CV8000 with a 40x/1.0NA water immersion objective lens, 50-μm pinhole, with three channels; 405 nm (Hoechst), 488 nm (GAP43), and 660 nm (β-tubulin III). Laser power for each channel was adjusted to ensure optimal signal-to-noise ratios, and 48 fields were imaged per well.

### Image Processing and Analysis

The open-source software CellProfiler (v4.2.1) was used for image QC, feature extraction, and analysis on a Windows-based workstation with dual Intel Xeon Platinum 8173M processors and 192GB RAM. Raw images were run through a pipeline set to calculate blurriness parameters. Images were analyzed in CellProfiler Analyst (v3.0.4) to obtain threshold values for the set quality parameters. The upper and lower thresholds were saved for future analyses.

A cell segmentation and skeletonization pipeline was developed in CellProfiler. The pipeline automatically flagged images that failed to pass QC. Primary object segmentation was performed using Hoechst 3342 stain of the nucleus. Secondary objects were identified using GAP43 staining of the soma by Otsu thresholding. As an additional aim of this pipeline was to study neuronal branching, β-tubulin III was used in the *Enhance or Suppress Features* module to enhance neurite structures using the Tubeness method and the smoothing scale set to 5. A new output image was saved as Neurite-enhances. Threshold module was used to reassign the pixel values that were below zero to zero and those above 1 to 1. A new output image was saved as Threshold_Neurite.

A Neurite-enhanced image was used to identify a secondary object, named Neurite, with Nucleus as the primary object using Otsu thresholding. Using the *Convert Objects to Image* module, the Neurite secondary objects were converted into single pixel wide image format and saved as Neurite_image. These images were used as input in the *Morphological Skeleton* module. The resulting skeletonized image was saved as Neurite_skeleton. A *Measure Object Skeleton* module was set to capture information regarding the number of axon and dendrite trunks, per branching structure, and the number of branches along the axons and dendrites. Nuclei was set to the seed object and Neurite_skeleton was set as the skeletonized image. Fill small holes was set to Ye, to avoid any false branch data points. The data was exported using *Export to Spreadsheet* module and saved in .csv format. A metadata file was prepared for each plate processed. A summary of the CellProfiler pipeline is described in **Figure S1**.

### Statistical and Benchmark Concentration Analysis

A total of 182,963 cells were imaged and analyzed across all time points and exposure conditions, based on Hoechst segmentation. Average values for each measure of each replicate were obtained by taking the median value per object and aggregating data by replicate, thus data represents the median value for a given measure across a given well. Comparisons between each experimental group and control was performed using a two-sided t-test in R (v4.3.3). Statistical significance was accepted with p < 0.05.

Benchmark concentrations (BMCs) and best fit BMC models were identified according to best practices for concentration-response modeling using BMDExpress v3.0, a free software package available through NIEHS, EPA, Health Canada, and Sciome (Yang et al., 2007). Original, untransformed data for each time point was uploaded and pre-filtered using One-Way ANOVA, with significance thresholded at p < 0.05 and fold change (FC) > |2| to identify measures with significance increasing or decreasing concentration response. Filtered data were modeled in BMDExpress with exponential (2, 3, 4, 5), linear, polynomial (2°, 3°, 4°), Hill, and power, and the best fit model was chosen based on the lowest Akaike information criteria (AIC). The benchmark response (BMR) type was set to 1 standard deviation relative to control with a confidence level of 0.95. Best fit models were chosen for each measure using nested chi square. Hill models were flagged if the ‘k’ parameter was 1/3 of the lowest possible dose. Benchmark concentrations (BMCs) were estimated from the Pb concentration-response model at each time point for each morphological metric. Concentration-response fit was assessed via examination of the best fit Pb concentration-response curves for individual morphology measures at each time point. Best benchmark concentrations estimated by BMDExpress greater than 10μM Pb were excluded from this analysis, as they exceeded the concentration range utilized in this study.

## Results

### Differentiation More Impactful on Cell Number Than Lead Exposure

Cell number was totaled for each image and the number of cells per image was averaged across each well (n = 6 per exposure condition at each time point). There was a marked decrease in the number of cells per well (and image) for all exposure conditions, including control, on Days 12, 15, and 18, relative to Day 6. This is expected, as the splitting steps on Days 7 and 10 of the differentiation protocol are stressful for all cells and only a subset survive the process (**Table S2**). On Day 6, the average number of cells captured in an image was 118.5, with non-significant (p > 0.05) increases in cell number across doses. By Day 12, there was a significant increase in cell number at 10μM Pb exposure (mean = 87.57 cells/image, p = 0.002) Pb conditions, relative to control (mean = 17.3 cells/image). On Days 15 and 18, no Pb exposure condition significantly impacted cell number relative to the control condition (**Figure 3**).

**Figure 3:**
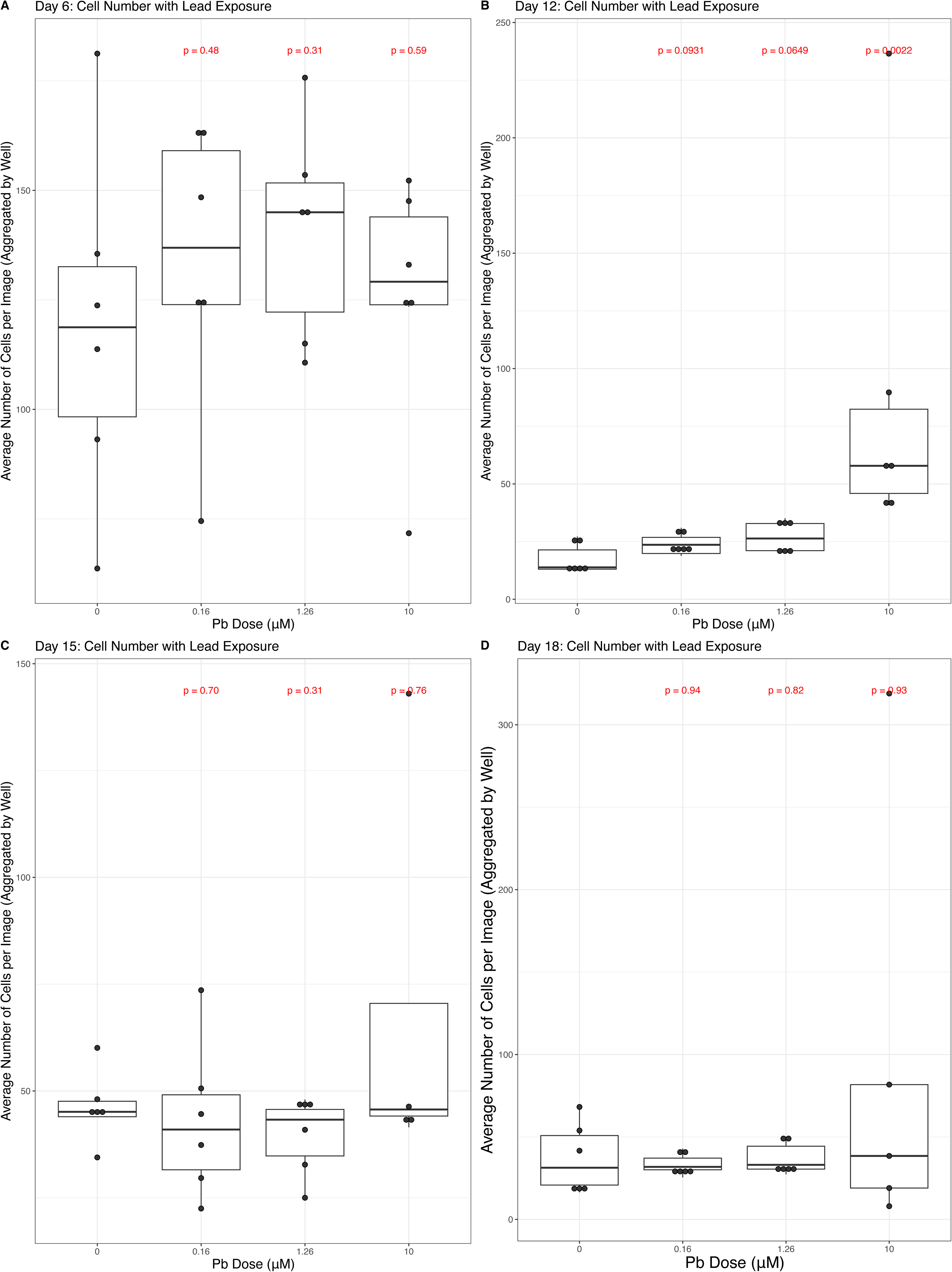
Cell Number and Lead (Pb). Average cell number was quantified per well across 48 fields per well. Boxplots describe the data, across all four exposure conditions (0uM, 0.16uM, 1.26uM, or 10uM Pb) on Days 6 (A), 12 (B), 15 (C), and 18 (D). Measures of statistical significance, determined using the Wilcoxon test, are indicated in red above each exposure condition, calculated relative to control.

### High Concentrations of Lead Exposure Impact Parameters of Cell Size

We next assessed parameters of cell size, including overall cell size as well as the size of the nucleus (**Figure 4**). Overall cell size was not impacted by either 0.16μM or 1.26μM Pb exposure, relative to control cells, on days 6, 12, and 15 (p > 0.05). By Day 18, the 1.26μM Pb condition resulted in a significant decrease in cell size (p = 0.002), relative to control cells. 10μM Pb exposure was associated with a significant decrease in overall cell size on Days 12 (p = 0.002, **Figure 4C)** and 15 (p = 0.009, **Figure 4E**), relative to control cells, with no significant difference detected on Days 6 (p > 0.05, **Figure 4A**) or 18 (p > 0.05, **Figure 4G**). Using BMDExpress, we estimated the best BMC estimate of the Pb concentration that significantly alters overall cell size and found significant best BMCs on Days 12 (1.85μM or 38.36μg/dL Pb) and 15 (8.99μM or 186.48μg/dL Pb, **Table S3**).

**Figure 4:**
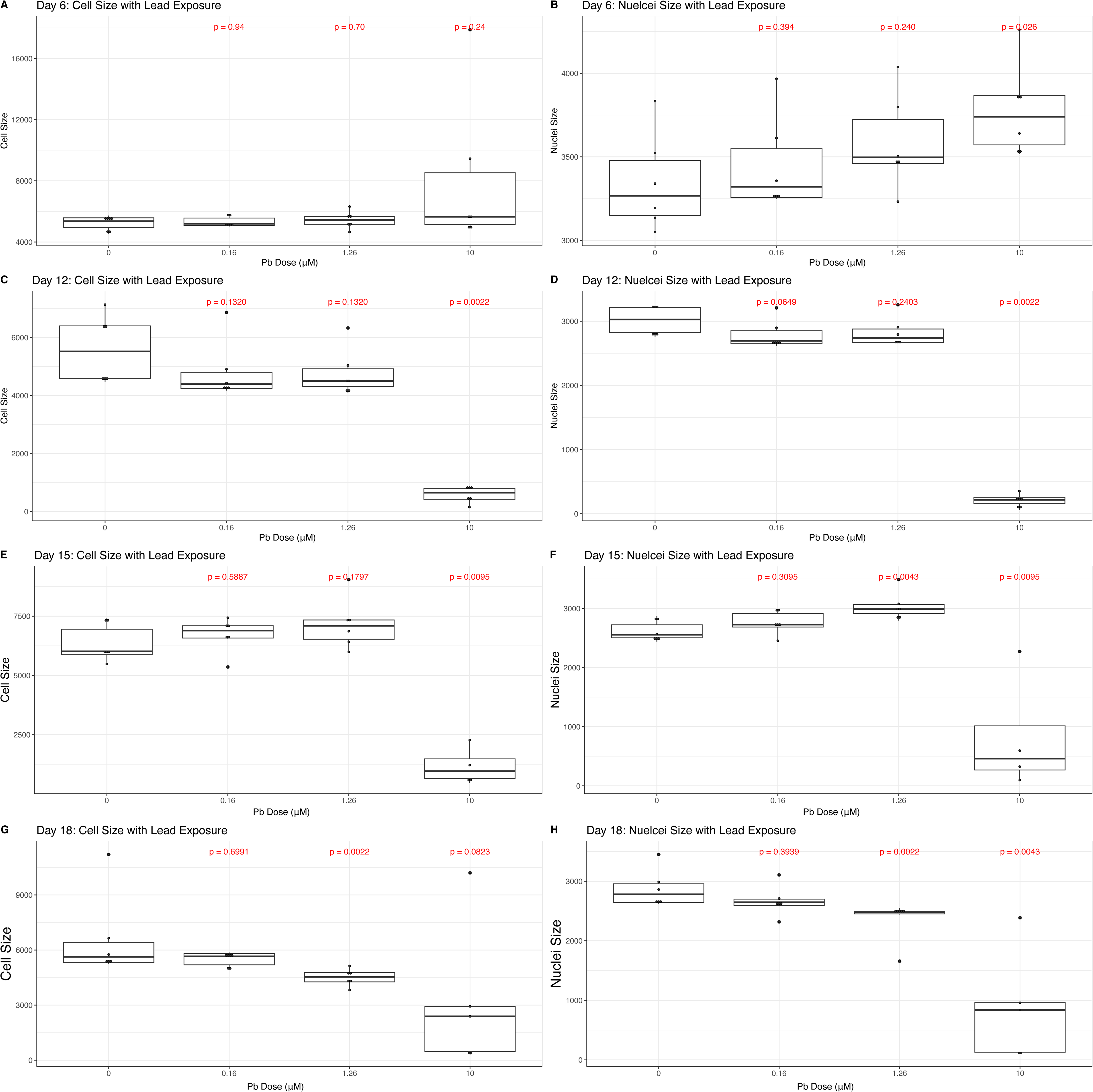
Cell Size and Lead (Pb). Average cell and nuclear size was quantified for each cell in CellProfiler, and averaged per well across 48 fields per well. Boxplots describe the data, across all four exposure conditions (0uM, 0.16uM, 1.26uM, or 10uM Pb) on Days 6 (A-B), 12 (C-D), 15 (E-F), and 18 (G-H). Measures of statistical significance, determined using the Wilcoxon test, are indicated in red above each exposure condition, calculated relative to control.

We also assessed impacts of Pb exposure on nuclear size throughout the time course. Beginning on Day 6, 10μM Pb exposure resulted in a significant increase in the size of the nucleus (p = 0.02, **Figure 4B**), and no changes seen with either of the lower concentrations. Nuclear size was reduced on Day 12 with 10μM Pb exposure, relative to control cells (p = 0.002), and again the lower concentrations had no significant effect (**Figure 4D**). On Day 12, BMDExpress estimated a best BMCs for alterations in nuclear size of 2.07μM Pb (42.98μg/dl Pb, **Table S3**). Nuclear size was more dynamic with Pb exposure during latter phases of SH-SY5Y differentiation. On Day 15, there was a significant increase in nuclei size with 1.26μM Pb exposure, relative to control (p = 0.004, **Figure 4F**), whereas on Day 18, this same exposure was associated with a decrease in nuclei size (p = 0.002, **Figure 4H**). 10μM Pb exposure continued to be associated with a significant decrease in the size of the nucleus at both of these time points (p < 0.05). On Day 15, the best BMC for nuclear size was 9.19μM Pb (190.53μg/dL Pb, **Table S3**) and this decreased notably on Day 18 to 2.58μM Pb (53.56μg/dL Pb, **Table S3**).

### Lead Exposure Perturbs the Expression of Neural Hallmarks During SH-SY5Y Differentiation

We quantified the intensity of two protein hallmarks of neural differentiation (β-tubulin III and GAP43) via fluorescently tagged antibodies (**Figure 5**) as well as the intensity Hoechst 33342 (**Figure S2**). There was no significant difference in the expression of β-tubulin III on Day 6, in any exposure condition relative to control (p > 0.05, **Figure 5A**). On Day 12, there was a significant increase in the expression of β-tubulin III with 0.16μM Pb exposure (p = 0.02) and a significant decrease in its expression in the 10μM Pb condition (p = 0.002, **Figure 5C**). By Day 12, BMDExpress predicted a best BMC for β-tubulin III of 1.19μM Pb (24.72μg/dL Pb, **Table S3**). On Day 15, expression of β-tubulin III continued to be significantly depressed in the 10μM Pb condition, relative to control (p = 0.009), with no significant effect seen in the 0.16μM or 1.26μM Pb conditions (**Figure 5E**). By Day 18, both 1.26μM (p = 0.008) and 10μM Pb. (p = 0.004) exposure resulted in a decrease in β-tubulin III expression, relative to control cells, with no significant change seen in the 0.16μM Pb condition (p = 0.06, **Figure 5G**). Day 15 and 18 had predicted best BMCs of 9.15μM Pb (189.6μg/dL Pb) and 1.15μM Pb (23.9μg.dL Pb, **Table S3**), respectively.

**Figure 5:**
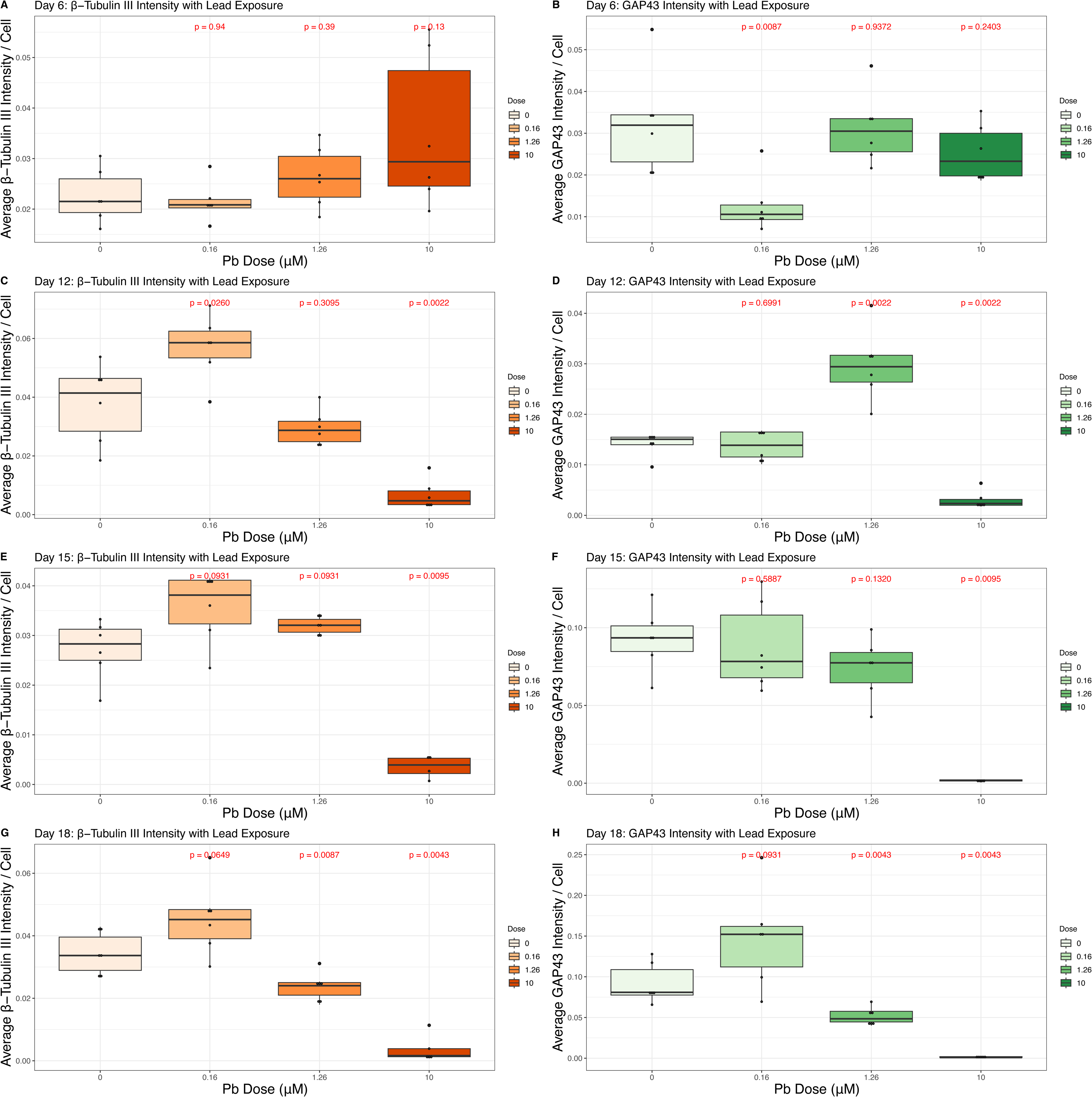
Neural Hallmarks and Lead (Pb). Immunofluorescent intensity of B-tubulin III and GAP43 were quantified in CellProfiler using the average intensity across all cells in a given image. Fluorescence intensity was averaged across all images to provide an aggregate measure per well. Boxplots describe the data, across all four exposure conditions (0uM, 0.16uM, 1.26uM, or 10uM Pb) on Days 6 (A-B), 12 (C-D), 15 (E-F), and 18 (G-H). Measures of statistical significance, determined using the Wilcoxon test, are indicated in red above each exposure condition, calculated relative to control.

The expression of GAP43 was affected by Pb exposure early in differentiation, with a significant decrease in its expression on Day 6 with 0.16μM Pb exposure (p = 0.008, **Figure 5B**). At Day 12, GAP43 expression was significantly increased with the 1.26μM Pb condition (p = 0.002) and significantly decreased in the 10μM Pb cells, relative to control (p = 0.002, **Figure 5D**). On Day 15, only the 10μM Pb condition significantly affected expression, with a decrease in GAP43 (p = 0.009, **Figure 5F**).

Interestingly, the estimated best BMC for GAP43 decreased substantially on Day 15 (2.23μM or 46.36μg/dL Pb) from the estimate on Day 12 of 9.59μM Pb (198.72μg/dL Pb, **Table S4**). As differentiation concluded on Day 18, both the 1.26μM (p = 0.004) and 10μM Pb (p = 0.004) conditions significantly depressed the expression of GAP43, relative to control cells (**Figure 5H**), with an estimated best BMC of 1.17μM Pb (24.4μg/dL Pb, **Table S4**) as determined by BMDExpress.

### Moderate and High Lead Exposure Decreases the Number and Length of Neural Projections

We used β-tubulin III staining, which is highly expressed in neurites (Latremoliere et al., 2018) to quantify the number and total length of neural projections from each cell during SH-SY5Y neural differentiation (**Figure 6**). Representative images of the skeletonization process used in this analysis in control cells can be found in **Figure S3**. Early in differentiation, on Day 6, Pb exposure had no effect on either the number or length of neural projections at any concentration, relative to control cells (p > 0.05, **Figure 6A-B**). By Days 12 and 15, 10μM Pb exposure alone resulted in a significant decrease in the number of neural projections (Day 12 p = 0.001 **Figure 6C**, Day 15 p = 0.01 **Figure 6E**) with between 0 and 1 projections per cell in the exposed condition compared to 2 to 3 projections in the control cells on these days. Best BMC estimates for number of projections per cell on Day 12 and 15 were 1.69μM Pb (35.18μg/dl Pb) and 5.45μM Pb (112.9μg/dL Pb), respectively (**Table S3**). The total length of projections also significantly decreased on Day 12 (p = 0.003, **Figure 6D** and Day 15 (p = 0.01 **Figure 6F**) in 10μM Pb exposed cells, relative to control cells. Best BMC estimates for total length of projections per cell on Day 12 and 15 were 1.76μM Pb (36.54μg/dl Pb) and 7.41μM Pb (153.59μg/dL Pb), respectively (**Table S3**). As differentiation concluded on Day 18, both 1.26μM and 10μM Pb exposure significantly decreased the number of neural projections (1.26μM Pb p = 0.03, 10μM Pb p = 0.01, **Figure 6G**) as well as their total length (1.26μM Pb p = 0.01, 10μM Pb p = 0.008, **Figure 6H**), relative to control cells. Best BMC estimates for number of projections per cell and total length of projections per cell were 2.01μM Pb (41.85μg/dL Pb) and 0.67μM Pb (13.83μg/dL Pb, **Table S3**), respectively.

**Figure 6:**
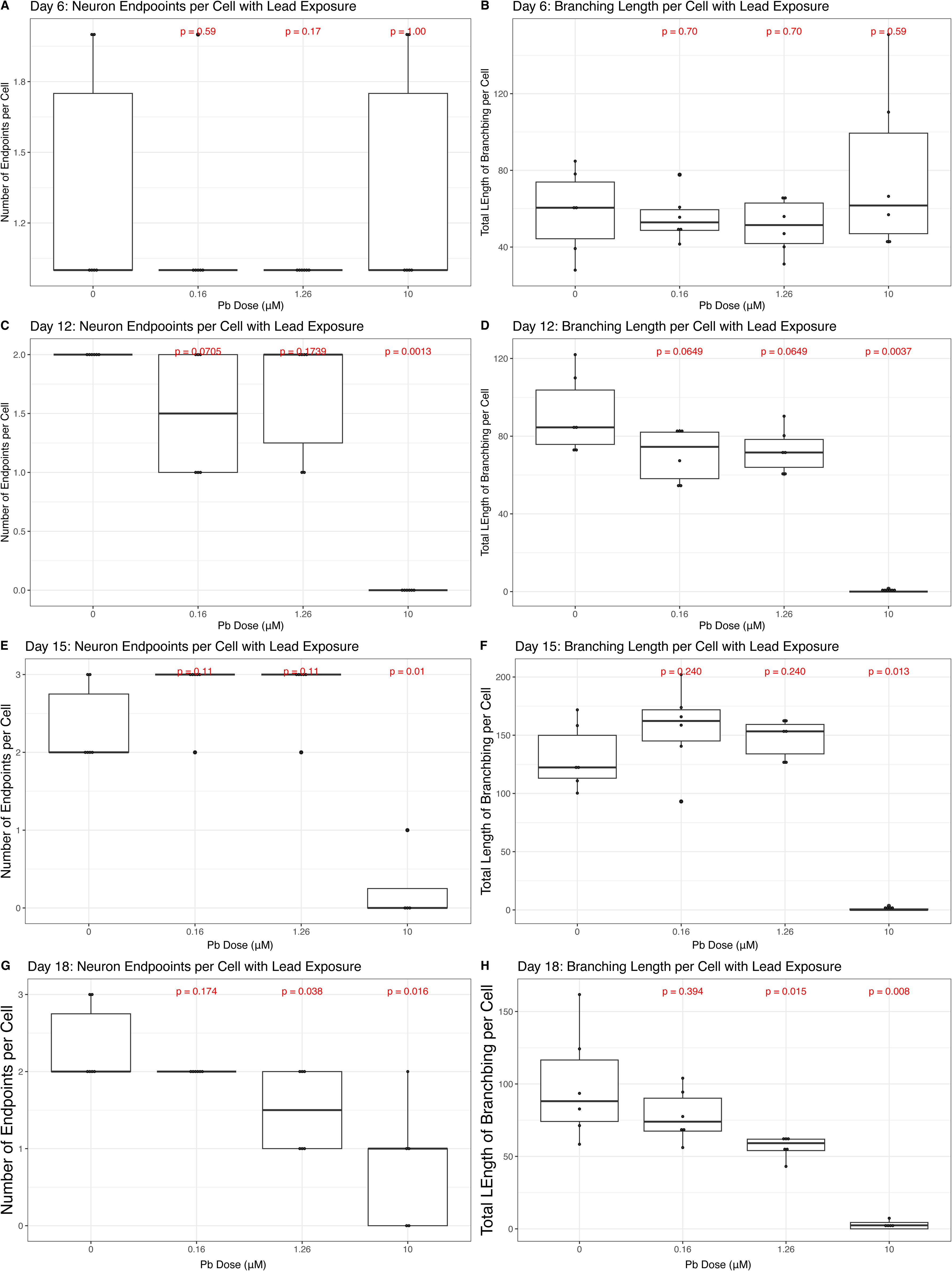
Neural Projections and Lead (Pb). The number and length of projections emanating from each cell was quantified using the skeletonization feature in CellProfiler. The average number of projections per cell and average total length of projections per cell were aggregated across all 48 images captured in each well. Boxplots describe the data, across all four exposure conditions (0uM, 0.16uM, 1.26uM, or 10uM Pb) on Days 6 (A-B), 12 (C-D), 15 (E-F), and 18 (G-H). Measures of statistical significance, determined using the Wilcoxon test, are indicated in red above each exposure condition, calculated relative to control.

## Discussion

We developed a high content imaging approach to assess the effects of environmentally relevant and continuous Pb exposure on cell morphology during the process of neural differentiation in the SH-SY5Y neuroblastoma cell model. We found Pb exposure to significantly affect the expression of both β-tubulin III and GAP43, as well as the formation and length of neurites, and that these effects were differentiation stage-specific and intensified with continued exposure.

### Lead Exposure Not Consistently Associated with Cell Number in Differentiating SH-SY5Y Cells

Pb exposure was not associated with increased cell number, during SH-SY5Y differentiation, with the exception of Day 12 when a significant increase was noted with the 1.26μM and 10μM Pb exposure conditions (**Figure 3**). Pb and other metals (e.g., zinc sulphate) have previously been shown to increase rates of cellular division and proliferation (Fathi & Farahzadi, 2018) and are thought to do so via the upregulation of cellular pathways related to stemness. Day 12 is a dynamic day for SH-SY5Y differentiation, corresponding to 24 hours after the addition of terminal differentiation factors including BDNF, and it may be that this state of heightened stemness may help explain the increased rate of cellular proliferation with Pb exposure. It is worth noting that this effect may be concentration-specific, as higher exposure levels have been shown to depress neural proliferation in related models (Körtje et al., 1991; Luo et al., 2023). It may also be that we would have observed a decrease in cellular proliferation and neurogenesis with continued Pb exposure had the terminal cells been kept in culture after differentiation concluded on Day 18, as others have seen with Pb exposure and a terminal neural cells (J. Li et al., 2024; Wang et al., 2013). This effect of increased proliferation during periods of stemness may also be specific to metals, as other exposure types (e.g., paraquat) result in the reduction of cellular proliferation (Colle et al., 2018).

### Lead Exposure Affects Parameters of Cell Size Differently Depending on Differentiation Stage

We evaluated total cell size as well as that of the nucleus, in relation to Pb concentration during SH-SY5Y differentiation (**Figure 5**). Total cell size was only impacted by 10μM Pb exposure on Days 12 and 15, exposure associated with a decrease in cell size. A major regulator of neural cell growth is the mechanistic target of rapamycin (mTOR) cascade (Crino, 2016) and Pb exposure has previously been associated with the decreased expression of mTOR genes in the hippocampus (Zhang et al., 2023). The mTOR pathway has a dynamic relationship with oxidative stress, a major cellular outcome of Pb exposure, as lower concentrations of reactive oxygen species have been shown to activate protein complexes within the mTOR cascade (Meijles et al., 2019), while higher concentrations repress these same complexes (M. Li et al., 2010). It may be that the low and moderate concentrations utilized here induced ROS concentrations capable of activating mTOR pathways enough to maintain cell size, while the 10μM Pb concentration produced a concentration of ROS that surpassed what is activating to the mTOR cascade, resulting in its suppression and a decrease in cell size. Future studies could investigate this intersection between Pb exposure, mTOR activation, and alterations in cell size throughout neurogenesis.

Counter to what was observed with whole cell, nuclear size was sporadically found to be elevated with Pb exposure, specifically on Day 6 (10μM Pb exposure) and Day 15 (1.26μM Pb exposure). Pb and other metals have been associated with disruptions to the nuclear membrane (Banfalvi et al., 2012) and chromatin condensation (Hernández-Ochoa et al., 2006), and Pb increases the expression of proteins related to neural nucleus development in the hippocampus (Fan et al., 2023). To our knowledge, this is the first report of nuclear size with Pb exposure during neural differentiation, and it may be that some of the changes here are due in part to aberrations in cell cycling. Nuclear size is known to fluctuate during interphase in preparation for cell division (Balachandra et al., 2022), and Pb exposure is thought to disrupt cell cycling through DNA damage (Yedjou et al., 2015). However, it is worth noting that nuclear expansion during cell cycling is thought to be proportional to an increase in overall cell size, and we did not observe a strong parallel association in this metric here, suggesting the observed effects on the nucleus were independent of Pb-induced changes in cell cycling and another mechanism of action may be at work. Additional work examining the effects of Pb exposure on subcellular compartments during distinct stages of cell cycling would help elucidate whether Pb effects on cell cycling play a role in the result seen here.

### Effect of Lead Exposure on Markers of Neural Differentiation is Stage-Specific

β-tubulin III is a microtubule cytoskeletal protein encoded by the human gene *TUBB3* and is predominantly expressed in neural cells (Radwitz et al., 2022). We observed greater variability in β-tubulin III expression in 10μM Pb exposed cells on Day 6 and a non-monotonic relationship with Pb exposure on Day 12, with 0.16μM Pb exposure increasing β-tubulin III expression, no change relative to control observed with 1.26μM Pb exposure, and a significant decrease in β-tubulin III expression with 10μM Pb exposure. Other exposures (e.g., carbon monoxide) (Dreyer-Andersen et al., 2018) and the induction of oxidative stress (Pérez Estrada et al., 2014) have been associated with an elevation of β-tubulin III during early differentiation in neural stem cell models, supporting the idea that cellular stress during periods of differentiation drive the upregulation of stem cell pathways. As differentiation progressed, this non-monotonic relationship disappeared, as both the moderate (1.26μM Pb) and high (10μM Pb) exposure conditions significantly decreased β-tubulin III expression by Day 18. Pb exposure in a static neural stem cell model (N2A) contributes to depressed cytoskeletal development and organization (Chen et al., 2022). However, it should be noted that this work employed a higher Pb exposure range (up to 50μM Pb, with the majority of effects seen above 25μM Pb) and examined effects in stable, non-differentiating cells. It is possible that the effects seen here are unique to the differentiation period and that cytoskeletal development may initially increase in the presence of Pb, while additional pathways promoting its expansion are also active, and that upon the conclusion of differentiation, sustained Pb exposure compromises what has been established. It would be prudent to expand this work in the future, employing multiple models of exposure timing, to assess how cytoskeletal structures adapt to exposures that begin and end at varying times.

GAP43 is commonly referred to as a growth or plasticity protein and has been well documented during neural differentiation as essential for neuronal plasticity and axon regeneration (Kosik et al., 1988). Previous work has demonstrated a robust decrease in GAP43 in differentiating SH-SY5Ys exposed to 5μM Pb for 24 hours, and these results are thought to be related to Pb-induced decreases in cellular calcium concentrations (Ayyalasomayajula et al., 2020). We also observed a decrease in GAP43 expression after 24 hours of Pb exposure (Day 6), though this effect was only seen at 0.16μM Pb exposure and not with either of the higher concentrations. We observed a recovery in GAP43 expression by Day 12 with 0.16μM Pb exposure, as expression was indistinguishable from control cells, and overexpression of this protein with the moderate dose of 1.26μM Pb. It is possible that the low and moderate Pb concentrations employed in this study induced increased expression of GAP43 during early differentiation because they were high enough to trigger pathways related to cellular plasticity and resilience, thus increasing GAP43 activity, as documented with other stressors (Dayem et al., 2014; Santos et al., 2015) but not so high as to trigger cytotoxic effects (as seen at the highest concentration of 10μM Pb). As with β-tubulin III, Pb exposure was associated with the depression of GAP43 expression as differentiation concluded, with significant effects seen at the moderate and high concentrations by Day 18, suggesting that sustained Pb exposure, beyond periods of stemness, impairs neuron development. This same relationship has been quantified with other metals in the SH-SY5Y model, including cadmium (Pak et al., 2014) and mercury (Chan et al., 2017), suggesting there may be a common mode of action on neurogenesis.

### Lead Exposure Decreases Morphological Signatures of Neural Differentiation

We found Pb exposure to have no effect on the number of or total length of neural projections during early differentiation (Day 6) and that only the highest exposure condition (10μM Pb) affected these metrics during the middle time points of Days 12 and 15. By Day 18, we observed a concentration-dependent decrease in both the number of axons and dendrites per cell and their combined length, with significant decreases with 1.26μM and 10μM Pb exposure. It is possible that this effect may have intensified had the cells been maintained in culture with Pb once differentiation had completed, as other metals have been shown to disrupt neurite outgrowth with sustained exposure in both *in vitro* and *in vivo* models (Chan et al., 2017; Feng et al., 2019). β-tubulin III expression also decreased in a dose-dependent manner on Day 18, with significant effects seen with 1.26μM and 10μM Pb exposure. As β-tubulin III is a major cytoskeletal protein in neural projections (Ferreira & Caceres, 1992) and its use here to quantify neural projection development likely explains the observed correlating results of decreased β-tubulin III expression and a decrease in neural projections under these conditions. It may be that had we quantified other cytoskeletal proteins, such as alpha tubulins or intermediate filaments (e.g, Types III, IV, and VI), which are integral components to the neural cytoskeletal framework (Hausrat et al., 2021; Kirkcaldie & Dwyer, 2017), we would have seen protein-specific effects. Indeed, Pb exposure depresses the expression of another neural structural protein, MAP2 (Ge et al., 2018) and alters the expression and function of phosphatases, which are key regulators of cytoskeletal proteins (Rahman et al., 2011). Interestingly, Pb has also been implicated in hyperphosphorylation of Tau (Gąssowska et al., 2016) and this molecular alteration to Tau has been well characterized in the generation of neurofibrillary tangles (Bussian et al., 2018), a major indicator of neurodegenerative disease (Lee et al., 2001).

### Public Health Relevance of Benchmark Concentration Estimates

The expression of proteins important to the success of neural differentiation (β-tubulin III and GAP43), as well as the morphology and development of neural projections were all affected by Pb exposure in this study. While not all measures were significantly impacted by the lower two doses employed here (0.16μM and 1.26μM Pb), the 10μM Pb dose regularly affected these measures during the course of SH-SY5Y differentiation into neuron-like cells. Our companion analysis of best BMC illustrated that important parameters of neural differentiation are likely affected by concentrations of Pb exposure that are currently (Neuwirth, 2018) and/or historically relevant (McFarland et al., 2022). This was particularly true in the case of the Day 18 measures, as all significant best BMCs fell between 0.66 and 2.58μM Pb (13.83μg/dL and 53.56μg/dL Pb). These concentrations correlate with BLLs that were relatively common in the US around the middle of the 20^th^ century (McFarland et al., 2022), when blood Pb monitoring was becoming more common, but Pb had not yet been phased out of commonly used products. This suggests that individuals who were children during these decades were exposed to levels of Pb that may have significantly perturbed neural differentiation and neurodevelopment. Continued work investigating whether Pb-induced changes to neural morphology during essential processes such as neural differentiation are warranted to expand our understanding of how developmental exposure to Pb may contribute to the lifetime risk of developing neurodegenerative disease (Wen et al., 2022). It is also important to note that these best BMCs correlate to BLLs that are still commonly seen in the US, in the case of childhood Pb poisoning from contaminated water or deteriorating housing (Levin et al., 2008), as well as around the world (Larsen & Sánchez-Triana, 2023). These results highlight the importance of eliminating Pb exposure in all instances, as these exposures are predicted to have significant repercussions for neurodevelopment.

### Limitations

The work described here relies on the SH-SY5Y neuroblastoma cell model, which is an imperfect model of neural differentiation as it does not represent the complex and dynamic environment of the brain during neurodevelopment. SH-SY5Y cells are also cancerous in origin, which influences their differentiation as well as their response to toxicants, as quantified by their viability and metabolism. The use of non-cancerous pluripotent cell lines from a range of donors and protocols intended to differentiate cells into multiple cell types in future experiments would help address these limitations.

### Summary and Public Health Significance

Our data demonstrates that continuous Pb exposure during neural differentiation has a significant impact on the expression of proteins key to this neurodevelopmental process, as well as cell morphology and the development of neural projections. Many of these effects were found to be differentiation stage-specific and estimated to be significant at relatively low exposure levels. These results support the hypothesis that environmental toxicants such as Pb perturb normal neuronal development and contribute to compromised cognitive outcomes.

## References

Ayyalasomayajula, N., Bandaru, M., Dixit, P. K., Ajumeera, R., Chetty, C. S., & Challa, S. (2020). Inactivation of GAP-43 due to the depletion of cellular calcium by the Pb and amyloid peptide induced toxicity: An in vitro approach. Chemico-Biological Interactions, 316, 108927. 10.1016/j.cbi.2019.108927

Balachandra, S., Sarkar, S., & Amodeo, A. A. (2022). The Nuclear-to-Cytoplasmic Ratio: Coupling DNA Content to Cell Size, Cell Cycle, and Biosynthetic Capacity. Annual Review of Genetics, 56, 165–185. 10.1146/annurev-genet-080320-030537

Banfalvi, G., Sarvari, A., & Nagy, G. (2012). Chromatin changes induced by Pb and Cd in human cells. Toxicology in Vitro: An International Journal Published in Association with BIBRA, 26(6), 1064– 1071. 10.1016/j.tiv.2012.03.016

Batool, S., Raza, H., Zaidi, J., Riaz, S., Hasan, S., & Syed, N. I. (2019). Synapse formation: From cellular and molecular mechanisms to neurodevelopmental and neurodegenerative disorders. Journal of Neurophysiology, 121(4), 1381–1397. 10.1152/jn.00833.2018

Bussian, T. J., Aziz, A., Meyer, C. F., Swenson, B. L., van Deursen, J. M., & Baker, D. J. (2018). Clearance of senescent glial cells prevents tau-dependent pathology and cognitive decline. Nature, 562(7728), 578–582. 10.1038/s41586-018-0543-y

Calogero, A. M., Mazzetti, S., Pezzoli, G., & Cappelletti, G. (2019). Neuronal microtubules and proteins linked to Parkinson’s disease: A relevant interaction? Biological Chemistry, 400(9), 1099–1112. 10.1515/hsz-2019-0142

Chan, M. C., Bautista, E., Alvarado-Cruz, I., Quintanilla-Vega, B., & Segovia, J. (2017). Inorganic mercury prevents the differentiation of SH-SY5Y cells: Amyloid precursor protein, microtubule associated proteins and ROS as potential targets. Journal of Trace Elements in Medicine and Biology, 41, 119–128. 10.1016/j.jtemb.2017.02.002

Chazotte, B. (2011). Labeling nuclear DNA with hoechst 33342. Cold Spring Harbor Protocols, 2011(1), pdb.prot5557. 10.1101/pdb.prot5557

Chen, L., Liu, Y., Jia, P., Zhang, H., Yin, Z., Hu, D., Ning, H., & Ge, Y. (2022). Acute lead acetate induces neurotoxicity through decreased synaptic plasticity-related protein expression and disordered dendritic formation in nerve cells. Environmental Science and Pollution Research International, 29(39), 58927–58935. 10.1007/s11356-022-20051-1

Colle, D., Farina, M., Ceccatelli, S., & Raciti, M. (2018). Paraquat and Maneb Exposure Alters Rat Neural Stem Cell Proliferation by Inducing Oxidative Stress: New Insights on Pesticide-Induced Neurodevelopmental Toxicity. Neurotoxicity Research, 34(4), 820–833. 10.1007/s12640-018-9916-0

Crino, P. B. (2016). The mTOR signalling cascade: Paving new roads to cure neurological disease. Nature Reviews. Neurology, 12(7), 379–392. 10.1038/nrneurol.2016.81

Dayem, A. A., Kim, B., Gurunathan, S., Choi, H. Y., Yang, G., Saha, S. K., Han, D., Han, J., Kim, K., Kim, J.-H., & Cho, S.-G. (2014). Biologically synthesized silver nanoparticles induce neuronal differentiation of SH-SY5Y cells via modulation of reactive oxygen species, phosphatases, and kinase signaling pathways. Biotechnology Journal, 9(7), 934–943. 10.1002/biot.201300555

Dreyer-Andersen, N., Almeida, A. S., Jensen, P., Kamand, M., Okarmus, J., Rosenberg, T., Friis, S. D., Martínez Serrano, A., Blaabjerg, M., Kristensen, B. W., Skrydstrup, T., Gramsbergen, J. B., Vieira, H. L. A., & Meyer, M. (2018). Intermittent, low dose carbon monoxide exposure enhances survival and dopaminergic differentiation of human neural stem cells. PloS One, 13(1), e0191207. 10.1371/journal.pone.0191207

Fan, S., Weixuan, W., Han, H., Liansheng, Z., Gang, L., Jierui, W., & Yanshu, Z. (2023). Role of NF-κB in lead exposure-induced activation of astrocytes based on bioinformatics analysis of hippocampal proteomics. Chemico-Biological Interactions, 370, 110310. 10.1016/j.cbi.2022.110310

Fathi, E., & Farahzadi, R. (2018). Zinc Sulphate Mediates the Stimulation of Cell Proliferation of Rat Adipose Tissue-Derived Mesenchymal Stem Cells Under High Intensity of EMF Exposure. Biological Trace Element Research, 184(2), 529–535. 10.1007/s12011-017-1199-4

Feng, J., Chen, S., Wang, Y., Liu, Q., Yang, M., Li, X., Nie, C., Qin, J., Chen, H., Yuan, X., Huang, Y., & Zhang, Q. (2019). Maternal exposure to cadmium impairs cognitive development of male offspring by targeting the Coronin-1a signaling pathway. Chemosphere, 225, 765–774. 10.1016/j.chemosphere.2019.03.094

Ferreira, A., & Caceres, A. (1992). Expression of the class III beta-tubulin isotype in developing neurons in culture. Journal of Neuroscience Research, 32(4), 516–529. 10.1002/jnr.490320407

Forscher, P., & Smith, S. J. (1988). Actions of cytochalasins on the organization of actin filaments and microtubules in a neuronal growth cone. The Journal of Cell Biology, 107(4), 1505–1516. 10.1083/jcb.107.4.1505

Gąssowska, M., Baranowska-Bosiacka, I., Moczydłowska, J., Tarnowski, M., Pilutin, A., Gutowska, I., Strużyńska, L., Chlubek, D., & Adamczyk, A. (2016). Perinatal exposure to lead (Pb) promotes Tau phosphorylation in the rat brain in a GSK-3β and CDK5 dependent manner: Relevance to neurological disorders. Toxicology, 347–349, 17–28. 10.1016/j.tox.2016.03.002

Ge, Y., Chen, L., Sun, X., Yin, Z., Song, X., Li, C., Liu, J., An, Z., Yang, X., & Ning, H. (2018). Lead-induced changes of cytoskeletal protein is involved in the pathological basis in mice brain. Environmental Science and Pollution Research International, 25(12), 11746–11753. 10.1007/s11356-018-1334-6

Glass, T. A., Bandeen-Roche, K., McAtee, M., Bolla, K., Todd, A. C., & Schwartz, B. S. (2009). Neighborhood psychosocial hazards and the association of cumulative lead dose with cognitive function in older adults. American Journal of Epidemiology, 169(6), 683–692. 10.1093/aje/kwn390

Gruart, A., Muñoz, M. D., & Delgado-García, J. M. (2006). Involvement of the CA3-CA1 synapse in the acquisition of associative learning in behaving mice. The Journal of Neuroscience: The Official Journal of the Society for Neuroscience, 26(4), 1077–1087. 10.1523/JNEUROSCI.2834-05.2006

Guo, J., Qiang, M., & Ludueña, R. F. (2011). The distribution of β-tubulin isotypes in cultured neurons from embryonic, newborn, and adult mouse brains. Brain Research, 1420, 8–18. 10.1016/j.brainres.2011.08.066

Hausrat, T. J., Radwitz, J., Lombino, F. L., Breiden, P., & Kneussel, M. (2021). Alpha- and beta-tubulin isotypes are differentially expressed during brain development. Developmental Neurobiology, 81(3), 333–350. 10.1002/dneu.22745

Hernández-Ochoa, I., Sánchez-Gutiérrez, M., Solís-Heredia, M. J., & Quintanilla-Vega, B. (2006). Spermatozoa nucleus takes up lead during the epididymal maturation altering chromatin condensation. Reproductive Toxicology (Elmsford, N.Y.), 21(2), 171–178. 10.1016/j.reprotox.2005.07.015

Hou, S., Yuan, L., Jin, P., Ding, B., Qin, N., Li, L., Liu, X., Wu, Z., Zhao, G., & Deng, Y. (2013). A clinical study of the effects of lead poisoning on the intelligence and neurobehavioral abilities of children. Theoretical Biology & Medical Modelling, 10, 13. 10.1186/1742-4682-10-13

Kirkcaldie, M. T. K., & Dwyer, S. T. (2017). The third wave: Intermediate filaments in the maturing nervous system. Molecular and Cellular Neurosciences, 84, 68–76. 10.1016/j.mcn.2017.05.010

Körtje, K. H., Körtje, D., & Rahmann, H. (1991). The application of energy-filtering electron microscopy for the cytochemical localization of Ca(2+)-ATPase activity in synaptic terminals. Journal of Microscopy, 162(Pt 1), 105–114. 10.1111/j.1365-2818.1991.tb03120.x

Kosik, K. S., Orecchio, L. D., Bruns, G. A., Benowitz, L. I., MacDonald, G. P., Cox, D. R., & Neve, R. L. (1988). Human GAP-43: Its deduced amino acid sequence and chromosomal localization in mouse and human. Neuron, 1(2), 127–132. 10.1016/0896-6273(88)90196-1

Larsen, B., & Sánchez-Triana, E. (2023). Global health burden and cost of lead exposure in children and adults: A health impact and economic modelling analysis. The Lancet Planetary Health, 7(10), e831–e840. 10.1016/S2542-5196(23)00166-3

Latremoliere, A., Cheng, L., DeLisle, M., Wu, C., Chew, S., Hutchinson, E. B., Sheridan, A., Alexandre, C., Latremoliere, F., Sheu, S.-H., Golidy, S., Omura, T., Huebner, E. A., Fan, Y., Whitman, M. C., Nguyen, E., Hermawan, C., Pierpaoli, C., Tischfield, M. A., … Engle, E. C. (2018). Neuronal-Specific TUBB3 Is Not Required for Normal Neuronal Function but Is Essential for Timely Axon Regeneration. Cell Reports, 24(7), 1865–1879.e9. 10.1016/j.celrep.2018.07.029

Lee, V. M., Goedert, M., & Trojanowski, J. Q. (2001). Neurodegenerative tauopathies. Annual Review of Neuroscience, 24, 1121–1159. 10.1146/annurev.neuro.24.1.1121

Levin, R., Brown, M. J., Kashtock, M. E., Jacobs, D. E., Whelan, E. A., Rodman, J., Schock, M. R., Padilla, A., & Sinks, T. (2008). Lead exposures in U.S. Children, 2008: Implications for prevention. Environmental Health Perspectives, 116(10), 1285–1293. 10.1289/ehp.11241

Levin, R., Zilli Vieira, C. L., Rosenbaum, M. H., Bischoff, K., Mordarski, D. C., & Brown, M. J. (2021). The urban lead (Pb) burden in humans, animals and the natural environment. Environmental Research, 193, 110377. 10.1016/j.envres.2020.110377

Li, H., Xue, X., Li, L., Li, Y., Wang, Y., Huang, T., Wang, Y., Meng, H., Pan, B., & Niu, Q. (2020). Aluminum-Induced Synaptic Plasticity Impairment via PI3K-Akt-mTOR Signaling Pathway. Neurotoxicity Research, 37(4), 996–1008. 10.1007/s12640-020-00165-5

Li, J., Hu, M., Liu, Y., Lu, R., & Feng, W. (2024). Lead exposure leads to premature neural differentiation via inhibiting Wnt signaling. Environmental Pollution (Barking, Essex: 1987), 363(Pt 2), 125232. 10.1016/j.envpol.2024.125232

Li, M., Zhao, L., Liu, J., Liu, A., Jia, C., Ma, D., Jiang, Y., & Bai, X. (2010). Multi-mechanisms are involved in reactive oxygen species regulation of mTORC1 signaling. Cellular Signalling, 22(10), 1469–1476. 10.1016/j.cellsig.2010.05.015

Luo, H., Li, J., Song, B., Zhang, B., Li, Y., Zhou, Z., & Chang, X. (2023). The binary combined toxicity of lithium, lead, and manganese on the proliferation of murine neural stem cells using two different models. Environmental Science and Pollution Research International, 30(2), 5047– 5058. 10.1007/s11356-022-22433-x

Markowitz, M. (2000). Lead poisoning. Pediatrics in Review, 21(10), 327–335. 10.1542/pir.21-10-327

McFarland, M. J., Hauer, M. E., & Reuben, A. (2022). Half of US population exposed to adverse lead levels in early childhood. Proceedings of the National Academy of Sciences, 119(11), e2118631119. 10.1073/pnas.2118631119

Meijles, D. N., Zoumpoulidou, G., Markou, T., Rostron, K. A., Patel, R., Lay, K., Handa, B. S., Wong, B., Sugden, P. H., & Clerk, A. (2019). The cardiomyocyte “redox rheostat”: Redox signalling via the AMPK-mTOR axis and regulation of gene and protein expression balancing survival and death. Journal of Molecular and Cellular Cardiology, 129, 118–129. 10.1016/j.yjmcc.2019.02.006

Mingaud, F., Mormede, C., Etchamendy, N., Mons, N., Niedergang, B., Wietrzych, M., Pallet, V., Jaffard, R., Krezel, W., Higueret, P., & Marighetto, A. (2008). Retinoid hyposignaling contributes to aging-related decline in hippocampal function in short-term/working memory organization and long-term declarative memory encoding in mice. The Journal of Neuroscience: The Official Journal of the Society for Neuroscience, 28(1), 279–291. 10.1523/JNEUROSCI.4065-07.2008

Mishra, R., Gupta, S. K., Meiri, K. F., Fong, M., Thostrup, P., Juncker, D., & Mani, S. (2008). GAP-43 is key to mitotic spindle control and centrosome-based polarization in neurons. Cell Cycle (Georgetown, Tex.), 7(3), 348–357. 10.4161/cc.7.3.5235

Morgan, R. K., Tapaswi, A., Polemi, K. M., Tolrud, E. C., Bakulski, K. M., Svoboda, L. K., Dolinoy, D. C., & Colacino, J. A. (2024). Environmentally Relevant Lead Exposure Impacts Gene Expression in SH-SY5Y Cells Throughout Neuronal Differentiation. bioRxiv: The Preprint Server for Biology.

Morgan, R. K., Wang, K., Svoboda, L. K., Rygiel, C. A., Lalancette, C., Cavalcante, R., Bartolomei, M. S., Prasasya, R., Neier, K., Perera, B. P. U., Jones, T. R., Colacino, J. A., Sartor, M. A., & Dolinoy, D. C. (2024). Effects of Developmental Lead and Phthalate Exposures on DNA Methylation in Adult Mouse Blood, Brain, and Liver: A Focus on Genomic Imprinting by Tissue and Sex. Environmental Health Perspectives, 132(6), 67003. 10.1289/EHP14074

Morita, S., & Miyata, S. (2013). Synaptic localization of growth-associated protein 43 in cultured hippocampal neurons during synaptogenesis. Cell Biochemistry and Function, 31(5), 400–411. 10.1002/cbf.2914

Neuwirth, L. S. (2018). Resurgent lead poisoning and renewed public attention towards environmental social justice issues: A review of current efforts and call to revitalize primary and secondary lead poisoning prevention for pregnant women, lactating mothers, and children within the U.S. International Journal of Occupational and Environmental Health, 24(3–4), 86–100. 10.1080/10773525.2018.1507291

Noctor, S. C., Martínez-Cerdeño, V., Ivic, L., & Kriegstein, A. R. (2004). Cortical neurons arise in symmetric and asymmetric division zones and migrate through specific phases. Nature Neuroscience, 7(2), 136–144. 10.1038/nn1172

Pak, E. J., Son, G. D., & Yoo, B. S. (2014). Cadmium inhibits neurite outgrowth in differentiating human SH-SY5Y neuroblastoma cells. International Journal of Toxicology, 33(5), 412–418. 10.1177/1091581814550338

Pérez Estrada, C., Covacu, R., Sankavaram, S. R., Svensson, M., & Brundin, L. (2014). Oxidative stress increases neurogenesis and oligodendrogenesis in adult neural progenitor cells. Stem Cells and Development, 23(19), 2311–2327. 10.1089/scd.2013.0452

Petroff, R. L., Cavalcante, R. G., Colacino, J. A., Goodrich, J. M., Jones, T. R., Lalancette, C., Morgan, R. K., Neier, K., Perera, B. P. U., Rygiel, C. A., Svoboda, L. K., Wang, K., Sartor, M. A., & Dolinoy, D. C. (2023). Developmental exposures to common environmental contaminants, DEHP and lead, alter adult brain and blood hydroxymethylation in mice. Frontiers in Cell and Developmental Biology, 11, 1198148. 10.3389/fcell.2023.1198148

Radwitz, J., Hausrat, T. J., Heisler, F. F., Janiesch, P. C., Pechmann, Y., Rübhausen, M., & Kneussel, M. (2022). Tubb3 expression levels are sensitive to neuronal activity changes and determine microtubule growth and kinesin-mediated transport. Cellular and Molecular Life Sciences: CMLS, 79(11), 575. 10.1007/s00018-022-04607-5

Rahman, A., Brew, B. J., & Guillemin, G. J. (2011). Lead dysregulates serine/threonine protein phosphatases in human neurons. Neurochemical Research, 36(2), 195–204. 10.1007/s11064-010-0300-6

Ribak, C. E., Korn, M. J., Shan, Z., & Obenaus, A. (2004). Dendritic growth cones and recurrent basal dendrites are typical features of newly generated dentate granule cells in the adult hippocampus. Brain Research, 1000(1–2), 195–199. 10.1016/j.brainres.2004.01.011

Ruckart, P. Z., Jones, R. L., Courtney, J. G., LeBlanc, T. T., Jackson, W., Karwowski, M. P., Cheng, P.-Y., Allwood, P., Svendsen, E. R., & Breysse, P. N. (2021). Update of the Blood Lead Reference Value—United States, 2021. MMWR. Morbidity and Mortality Weekly Report, 70(43), 1509–1512. 10.15585/mmwr.mm7043a4

Santos, N. A. G., Martins, N. M., Sisti, F. M., Fernandes, L. S., Ferreira, R. S., Queiroz, R. H. C., & Santos, A. C. (2015). The neuroprotection of cannabidiol against MPP+-induced toxicity in PC12 cells involves trkA receptors, upregulation of axonal and synaptic proteins, neuritogenesis, and might be relevant to Parkinson’s disease. Toxicology in Vitro, 30(1, Part B), 231–240. 10.1016/j.tiv.2015.11.004

Scortegagna, M., Chikhale, E., & Hanbauer, I. (1998). Effect of lead on cytoskeletal proteins expressed in E14 mesencephalic primary cultures. Neurochemistry International, 32(4), 353–359. 10.1016/s0197-0186(97)00101-0

Seri, B., García-Verdugo, J. M., McEwen, B. S., & Alvarez-Buylla, A. (2001). Astrocytes give rise to new neurons in the adult mammalian hippocampus. The Journal of Neuroscience: The Official Journal of the Society for Neuroscience, 21(18), 7153–7160. 10.1523/JNEUROSCI.21-18-07153.2001

Shen, Y., Mani, S., & Meiri, K. F. (2004). Failure to express GAP-43 leads to disruption of a multipotent precursor and inhibits astrocyte differentiation. Molecular and Cellular Neurosciences, 26(3), 390–405. 10.1016/j.mcn.2004.03.004

Shipley, M. M., Mangold, C. A., & Szpara, M. L. (2016). Differentiation of the SH-SY5Y Human Neuroblastoma Cell Line. Journal of Visualized ExperimentslJ: JoVE, 108. 10.3791/53193

Silver, J., Lorenz, S. E., Wahlsten, D., & Coughlin, J. (1982). Axonal guidance during development of the great cerebral commissures: Descriptive and experimental studies, in vivo, on the role of preformed glial pathways. The Journal of Comparative Neurology, 210(1), 10–29. 10.1002/cne.902100103

UNICEF. (2020, July 30). The toxic truth | UNICEF. https://www.unicef.org/reports/toxic-truth-childrens-exposure-to-lead-pollution-2020

Wang, B., Feng, G., Tang, C., Wang, L., Cheng, H., Zhang, Y., Ma, J., Shi, M., & Zhao, G. (2013). Ginsenoside Rd maintains adult neural stem cell proliferation during lead-impaired neurogenesis. Neurological Sciences: Official Journal of the Italian Neurological Society and of the Italian Society of Clinical Neurophysiology, 34(7), 1181–1188. 10.1007/s10072-012-1215-6

Wen, Q., Verheijen, M., Wittens, M. M. J., Czuryło, J., Engelborghs, S., Hauser, D., van Herwijnen, M. H. M., Lundh, T., Bergdahl, I. A., Kyrtopoulos, S. A., de Kok, T. M., Smeets, H. J. M., Briedé, J. J., & Krauskopf, J. (2022). Lead-exposure associated miRNAs in humans and Alzheimer’s disease as potential biomarkers of the disease and disease processes. Scientific Reports, 12(1), 15966. 10.1038/s41598-022-20305-5

White, R. F., Diamond, R., Proctor, S., Morey, C., & Hu, H. (1993). Residual cognitive deficits 50 years after lead poisoning during childhood. British Journal of Industrial Medicine, 50(7), 613–622. 10.1136/oem.50.7.613

WHO. (2020, June 1). 10 chemicals of public health concern. https://www.who.int/news-room/photo-story/photo-story-detail/10-chemicals-of-public-health-concern

Xie, J., Wu, S., Szadowski, H., Min, S., Yang, Y., Bowman, A. B., Rochet, J.-C., Freeman, J. L., & Yuan, C. (2023). Developmental Pb exposure increases AD risk via altered intracellular Ca2+ homeostasis in hiPSC-derived cortical neurons. The Journal of Biological Chemistry, 299(8), 105023. 10.1016/j.jbc.2023.105023

Yang, L., Allen, B. C., & Thomas, R. S. (2007). BMDExpress: A software tool for the benchmark dose analyses of genomic data. BMC Genomics, 8, 387. 10.1186/1471-2164-8-387

Yedjou, C. G., Tchounwou, H. M., & Tchounwou, P. B. (2015). DNA Damage, Cell Cycle Arrest, and Apoptosis Induction Caused by Lead in Human Leukemia Cells. International Journal of Environmental Research and Public Health, 13(1), ijerph13010056. 10.3390/ijerph13010056

Zhang, B., Li, H., Wang, Y., Li, Y., Zhou, Z., Hou, X., Zhang, X., & Liu, T. (2023). Mechanism of autophagy mediated by IGF-1 signaling pathway in the neurotoxicity of lead in pubertal rats. Ecotoxicology and Environmental Safety, 251, 114557. 10.1016/j.ecoenv.2023.114557

