## Supplementary Figure 1 for "Environmentally Relevant Lead Exposure Alters Cell Morphology and Expression of Neural Hallmarks During SH-SY5Y Neuronal Differentiation"

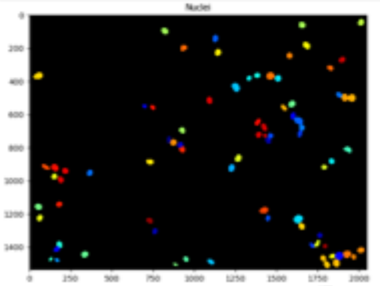


- Images
- Metadata
- Names and Types
- Groups
- Correct Illumination Apply
- **Identify Primary Objects**
- **Identify Primary Objects**

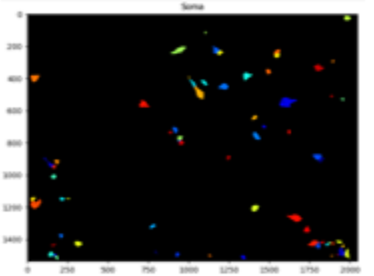


- Enhance or Suppress Features
- **Identify Secondary Objects**

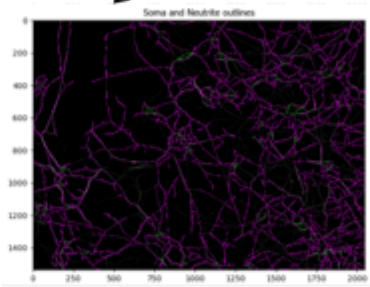


- Convert Objects To Image
- **Morphological Skeleton**

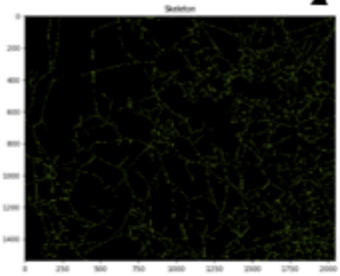


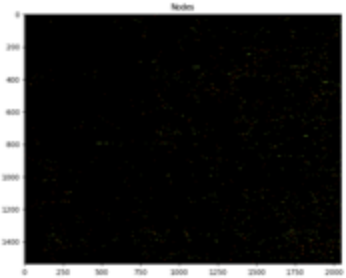


- Measure Image Skeleton
- Overlay Outlines
- Overlay Outlines
- Save Images
- Save Images
- Export to Database
- Export to Spreadsheet
