## Supplementary figures and images for "Environmentally Relevant Lead Exposure Alters Cell Morphology and Expression of Neural Hallmarks During SH-SY5Y Neuronal Differentiation"

### Supplemental Figure 3

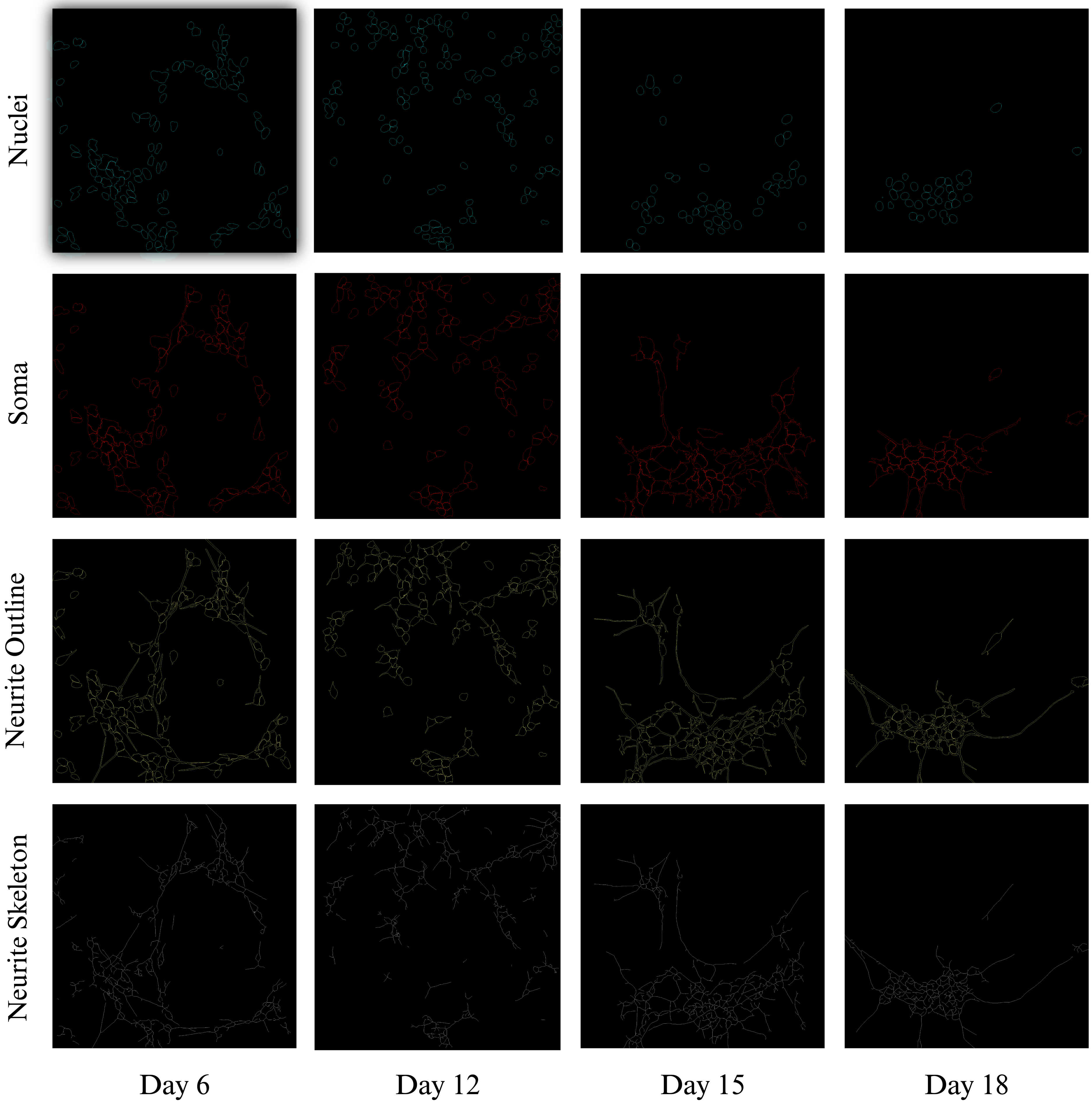

### Supplementary Figure 2

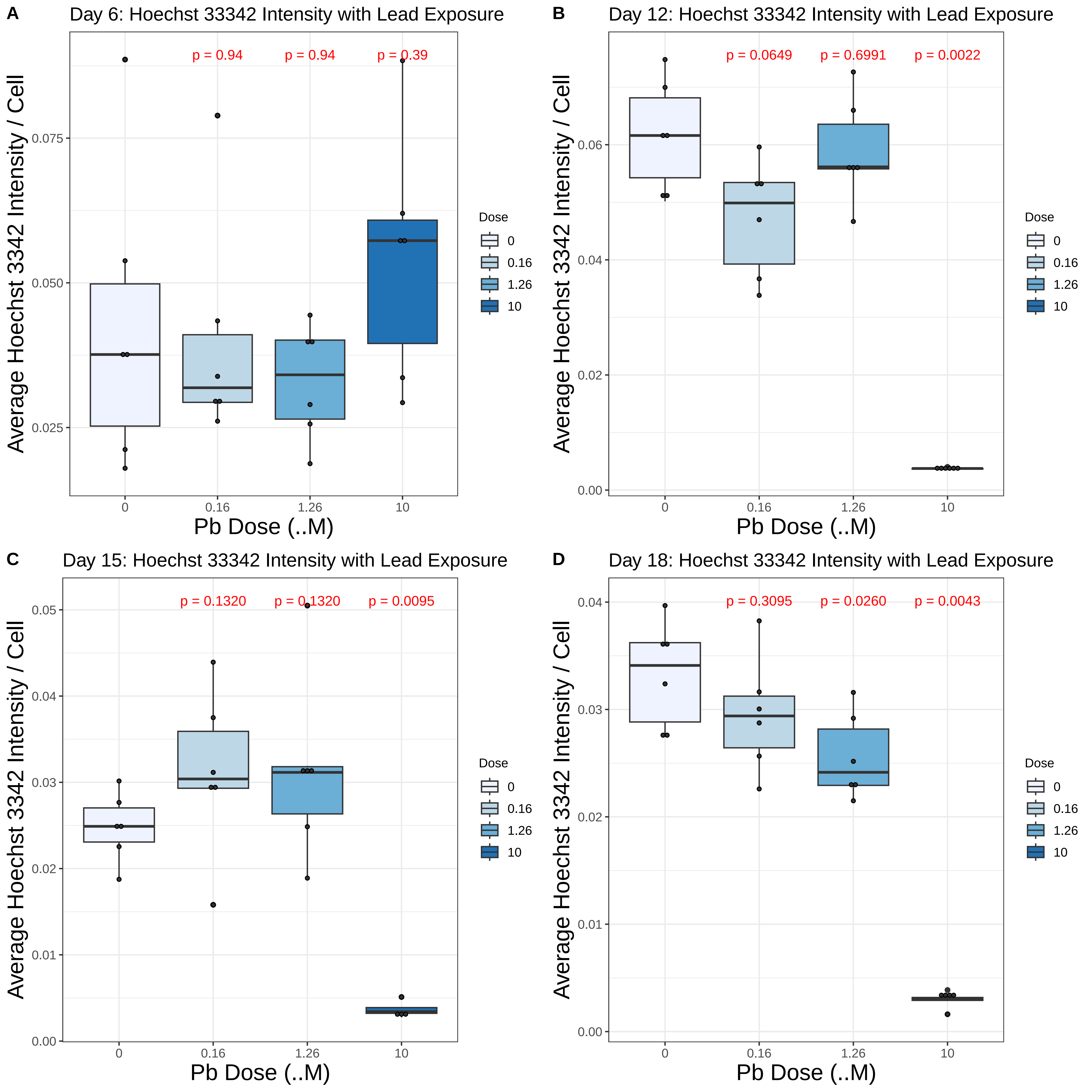
