## Supplemental File 1 for "Environmentally Relevant Lead Exposure Alters Cell Morphology and Expression of Neural Hallmarks During SH-SY5Y Neuronal Differentiation"

**SH-SY5Y Staining Protocol**

Materials Needed:

| - Cell line in culture: SH-SY5Y | - Antibodies/stains of interest: |
| --- | --- |
| - Media used for cells | - β tubulin III conjugate (Thermo, 50-4510-82)   - eFluor 660, host: mouse |
| - 16% paraformaldehyde (PFA) | - GAP43 Primary (Abcam, ab75810)   - Rabbit monoclonal |
| - PBS (pH 7.4) | - GAP43 Secondary (Abcam, ab150077)   - Fluor 488, host: goat, anti-rabbit |
| - Normal Goat Serum (NGS) | - Hoechst 33342 (Thermo, H3570) |
| - Tin foil/dark space to incubate |  |

Protocol:

A) Make PBS-T – you’ll need enough to rinse each well 10x
   a. Add Tween 20 to PBS at a 1:1000 concentration.
      i. 50mL PBS + 50µL Tween 20
B) Fix Cells with PFA – prepare and use in **chemical hood**
1. Dilute 16% to a 4% PFA solution (1:4) in media.
2. Gently remove media from cells.
3. Add enough 4% PFA solution to cover the bottom of each well (e.g., 100µL for a 24 well plate).
4. Incubate at room temperature for 20 minutes.
5. Remove 4% PFA solution and rinse each well with PBS-T one time.
C) Permeabilize cells with Triton X – prepare and use in **biosafety cabinet**
1. Add Triton X to PBS at a 1:1000 dilution.
2. Add enough Triton X-PBS solution to cover the bottom of each well.
3. Incubate at room temperature for 15 minutes.
4. Remove Triton X-PBS and rinse each well with PBS-T two times.
D) Primary Stain – work in the dark/protect antibodies from light as much as possible, prepare and use in **biosafety cabinet**
1. Create primary stain solution:
   a. For 1 24-well plate, 100µL per well:
      i. 2.5mL NGS
      ii. 1.25µL Hoechst 33342 (0.5µL/mL)
      iii. 16.5µL β-tubulin III conjugate antibody (6.6µL/mL)
      iv. 12.5µL GAP43 primary (5µL/mL)
2. Add enough primary stain solution to cover the bottom of each well.
3. Incubate 1 hour in the dark at room temperature.
4. Remove primary stain and rinse with PBS-T three times.
E) Secondary Stain – work in the dark/protect antibodies from light as much as possible, prepare and use in **biosafety cabinet**
1. Create secondary stain solution:
   a. For 1 24-well plate, 100µL per well:
      i. 2.5mL NGS
      ii. 7.5µL GAP43 secondary (3µL/mL)
2. Add enough primary stain solution to cover the bottom of each well.
3. Incubate 1 hour in the dark at room temperature.
4. Remove primary stain and rinse with PBS-T three times.
5. Add enough PBS-T back to each well to cover.
F) Imaging
1. Image on CV8000 using the following channels:
   a. 405 nm (Hoechst)
   b. 488 nm (GAP43)
   c. 660 nm (β-tubulin III)
